# The 3D ultrastructural morphology of a marine ammonia-oxidizing archaeon

**DOI:** 10.1101/2023.12.12.571259

**Authors:** Yangkai Zhou, An Yan, Jiawen Yang, Wei He, Shuai Guo, Yifan Li, Yanchao Dai, Xijiang Pan, Dongyu Cui, Olivier Pereira, Wenkai Teng, Ran Bi, Songze Chen, Lu Fan, Peiyi Wang, Yan Liao, Wei Qin, Sen-Fang Sui, Yuanqing Zhu, Chuanlun Zhang, Zheng Liu

## Abstract

*Nitrososphaerota*, formerly known as *Thaumarchaeota*, constitute a diverse and widespread group of ammonia-oxidizing archaea (AOA) inhabiting ubiquitously in marine and terrestrial environments, playing a pivotal role in global nitrogen cycling. Despite their significance in Earth’s ecosystems, the cellular organization of AOA remains largely unexplored, leading to a significant question unanswered of how the machinery of these organisms underpins metabolic functions. In this study, we combined chromatic-aberration-corrected cryo-electron tomography (cryo-ET), scanning transmission electron microscopy (STEM), and energy dispersive X-ray spectroscopy (EDS) to unveil the cellular organization and elemental composition of *Nitrosopumilus maritimus* SCM1, a representative member of marine *Nitrososphaerota*. Our tomograms show the native ultrastructural morphology of SCM1 and one to several dense storage granules in the cytoplasm. STEM-EDS analysis identifies two types of storage granules in which one type is possibly composed of polyphosphate, while the other type is polyhydroxyalkanoate. Collectively, these findings provide visual evidence for the resilience of AOA in the vast oligotrophic marine environment.

## Introduction

Comprehending the intricate cellular structure of microorganisms holds paramount importance in advancing our understanding of metabolic functions in ecology, evolution, and biogeochemistry (González-Pech et al., 2023). Despite remarkable advancements in cryo-electron microscopy (cryo-EM) technology and the potential value of archaeal structural biology in medical and industrial applications (Shin et al., 2014), the realm of structural biology of marine archaea remains relatively underexplored, necessitating efforts to comprehensively grasp their cellular structure and physiological mechanisms (Lipp et al., 2008; Stahl and de la Torre, 2012).

Ammonia-oxidizing archaea (AOA), belonging to the currently classified *Nitrososphaerota*, derive energy from the oxidation of ammonia to nitrite, initiating nitrification process that is vital in the aquatic nitrogen cycle (Stahl and de la Torre, 2012; Walker et al., 2010). The resulting nitrite, a byproduct of AOA activity, can be further transformed into nitrate by nitrite-oxidizing bacteria, and ultimately eliminated from the ecosystem as it converts into nitrogen gas through denitrifying bacteria (Martikainen, 2022). Notably, *Nitrosopumilus maritimus* SCM1 stands out as the first isolate of AOA (Konneke et al., 2005), exemplifying one of the most prevalent ammonia-oxidizing microbes in the oceans and actively driving global nitrogen cycling.

Previous studies have delved into various aspects of *Nitrosopumilus maritimus* SCM1, including its physiology like ammonium oxidation, stress adaptation, carbon fixation, genome, metabolome, lipidome, evolution and ecology (Abby et al., 2020; Hodgskiss et al., 2023; Kim et al., 2019; Kitzinger et al., 2020, 2019; Kraft et al., 2022; Law et al., 2021a, 2021b; Leavitt et al., 2023; Li et al., 2018; Martens-Habbena et al., 2009; Qin et al., 2020, 2018, 2014; Walker et al., 2010; Wan et al., 2023; Yang et al., 2021). However, our knowledge of its cellular structure remains limited. Within an archaeal cell, the cellular components are organized in a specific arrangement. Starting from the cell surface and moving inward, these components typically include an s-layer, pseudoperiplasmic space, cytoplasmic membrane, nucleoid, ribosomes. Additionally, certain archaeal cells may possess additional structures like vesicles, archaellum and storage granules (van Wolferen et al., 2022). Several studies have reported the presence of S-layer proteins on the surface of SCM1 cells, suggesting its existence as a cell wall (Qin et al., 2017; Urakawa et al., 2011). Additionally, high abundance of ribosomes numbering ∼1000 in each cell has been observed in SCM1 (Urakawa et al., 2011), which likely provide advantage in coping with extreme environmental conditions (Yin et al., 2018).

However, the cellular structure of AOA, including for example the presence and function of storage granules, remains inadequately understood due to limited research efforts. Storage granules play a pivotal role in microorganisms by enabling them to withstand fluctuations in nutrient availability. Although extensive investigations have been conducted on storage granules in certain bacterial, thermophilic archaea, and eukaryotic organisms (Gal et al., 2017; Sarkar et al., 2021; Tocheva et al., 2013; Toso et al., 2011, 2016; Ward et al., 2012), their existence and functionality in marine archaea, particularly in SCM1, have yet to be explored in detail (Qin et al., 2017; Urakawa et al., 2011). This knowledge gap necessitates further exploration to unravel the cellular structure and evaluate the presence and function of storage granules in marine archaea, and thereby expanding our understanding in this uncharted domain.

This study aimed to address the knowledge gap in structural biology of AOA by employing cutting-edge techniques that combine chromatic-aberration-corrected cryo-electron microscopy (cryo-EM), cryo-electron tomography (cryo-ET), scanning transmission electron microscopy (STEM), and energy-dispersive X-ray spectroscopy (EDS). These technologies allowed us to visualize the cell morphology of the AOA type strain *Nitrosopumilus maritimus* SCM1 at ultra-high resolution, which demonstrated the presence of two types of storage granules and cell division patterns. These findings provide new insight into the potential metabolic functions of storage granules in marine archaea, while shedding light on the high-resolution characteristics of SCM1 cells in energy storage. The use of chromatic-aberration-corrected cryo-ET, which was pioneered in this study, represents an innovative technology for effectively and comprehensively visualizing both archaeal and bacterial cells at submicron scales.

## Results

### Whole-cell cryo-electron tomography of *Nitrosopumilus maritimus* SCM1

We monitored cell growth in *Nitrosopumilus maritimus* SCM1 cultures by analyzing nitrite concentration. As shown in **Supplementary Figure 1**, during the nine days’ cultivation period, nitrite concentration continued rise in the first seven days. We harvested SCM1 cells between day 6 and day 8, corresponding to the middle to later exponential phase. The purity of the cultures was evaluated using the qPCR method, and the results are presented in **Supplementary Table 1**.

Initially, we examined the morphology of *Nitrosopumilus maritimus* SCM1 cells by traditional cryo-EM. Our observations revealed that SCM1 cells showed a typical rod shape, with diameters ranging from 320 to 410 nm, lengths spanning from 590 to 1350 nm and the mean length-diameter ratio around 2.43 ± 0.57 (**Supplementary Figure 2**). Subsequently, cryo-ET experiment elucidated the cellular organization of SCM1 cells and provided a native three-dimensional (3D) structural information of the cell. Given the considerable thickness of the SCM1 (averaged diameter ∼360nm, **Supplementary Figure 2**), the tilt-series image acquisition could not be completed. The thickness would double at ±60°, rendering it unsuitable for conventional cryo-EM data acquisition (**Supplementary Figure 3**). To overcome this difficulty, we employed the first chromatic-aberration-corrected cryo-EM in the world, of which can increase a signal/noise ratio for the thick specimens by corrected inelastic scattering. With the Cc corrector, images can be obtained at ±60°, and 3D tomogram can be reconstructed with less missing wedges. The 3D structure of native SCM1 revealed a highly organized proteinaceous surface layer (S-layer) enveloping the entire cell surface, and cell membrane exhibited well-defined, smooth, and continuous features (**Figure 1** and **Supplementary Movie 1**). Notably, an area rich in nucleic acids and dozens of ribosomes were observed in the cytoplasm of SCM1 (**Figure 1** and **Figure 2**) Interestingly, the number of ribosomes in SCM1 were ranged between 30-100 in each cell, which appeared to be significantly less than those found in eukaryotes (Xing et al., 2023) or other prokaryotes (Xue et al., 2022) that observed by cryo-ET, lending a support for the slow growth of SCM1 that typically observed in the laboratory culture (**Supplementary Figure 1**). Moreover, this result highlights the superior accuracy of cryo-ET in quantifying ribosomes at the three-dimensional level compared to counting them in two-dimensional microscopic images (Urakawa et al., 2011).

**Figure 1.**
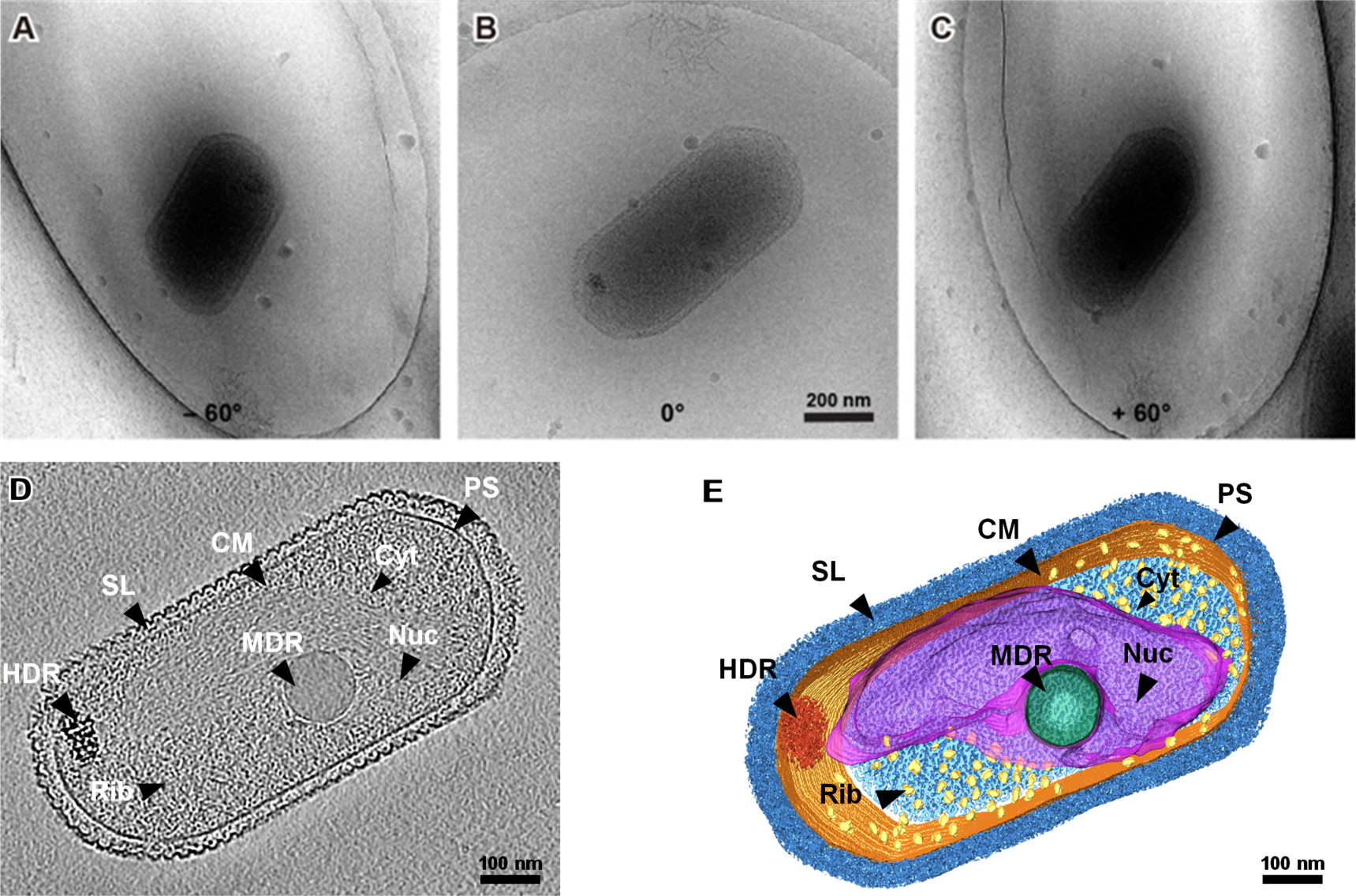
Whole-cell cryo-electron tomogram of *Nitrosopumilus maritimus* SCM1. (A-C) Cryo-EM images of SCM1 at – 60°, 0°, and + 60° angles. (D) Central section through a tomogram showed the cellular structure of SCM1. (E) 3D segmentation of SCM1 cell. MDR, medium-density region; HDR, high-density region; CM, cytoplasmic membrane; SL, surface layer; Nuc, nucleoid; Rib, ribosome; PS, pseudoperiplasmic space; and Cyt, cytoplasm, are displayed.

**Figure 2.**
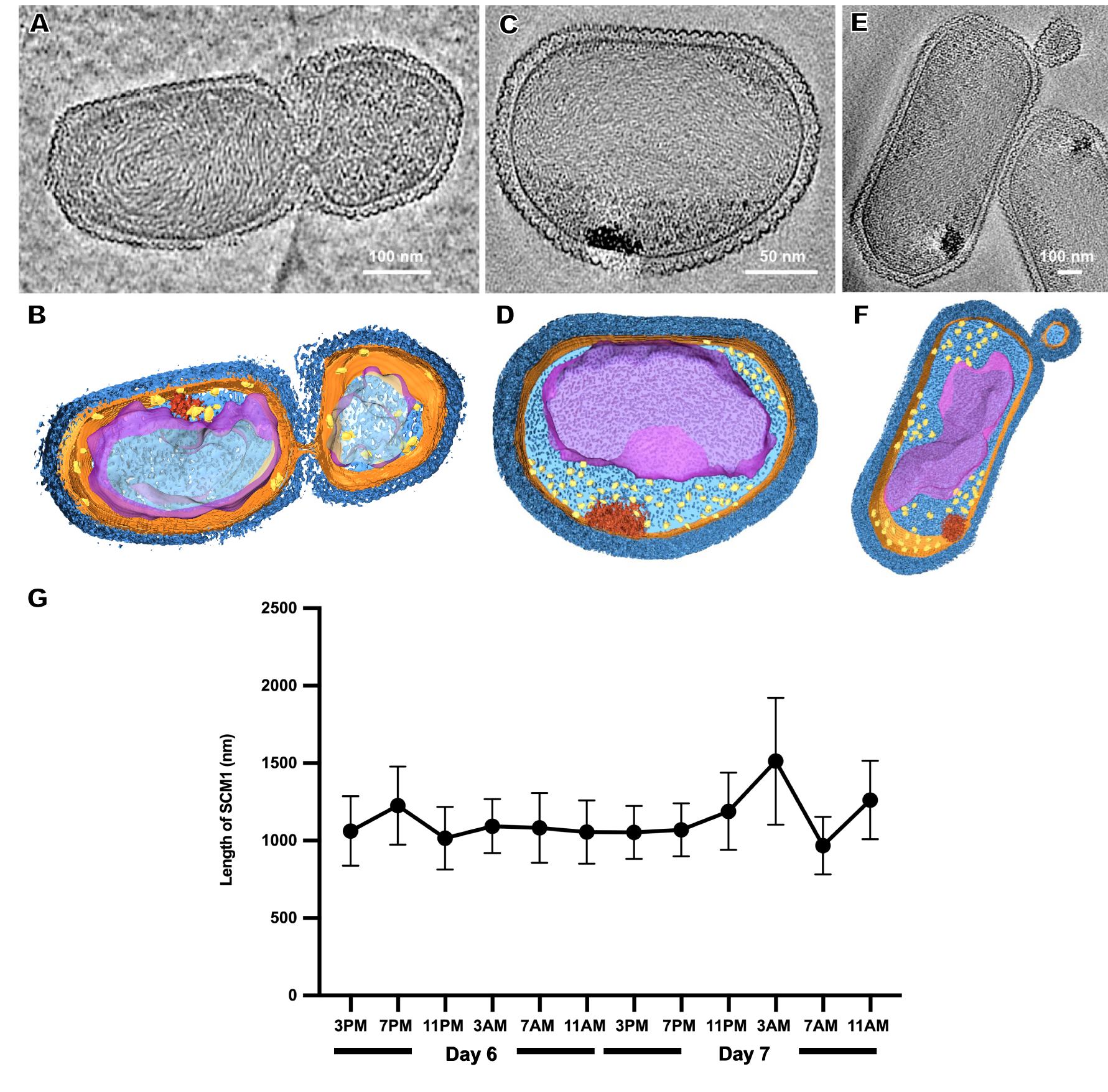
Observation of cell division in *Nitrosopumilus maritimus* SCM1. (A, C, E) Slices through tomograms of SCM1 in the exponential phase. (B, D, F) 3D segmentation of (A, C, E). S-layer (dark blue), cell membrane (orange); cytoplasm (light blue); nucleoid (purple); ribosomes (yellow); high-density region (red). (G) The average length of SCM1 at different time points. SCM1 cells were analyzed every 4 hours for 48 hours period in the late exponential phase (*n* = 40).

### Observation of cell division in *Nitrosopumilus maritimus* SCM1

We examined several cells in the exponential phase and identified instances of unequal divisions (**Figure 2A** and **2B**). The central region of SCM1 first exhibited constriction, followed by elongation towards two ends before eventually dividing into two cells. Notably, the cell division of SCM1 did not follow a homomorphic pattern, with the larger daughter cell assuming an oblate spheroidal shape, resembling the shape of another independently observed cell in **Figure 2C** and **2D**. Additionally, we observed a putative budding state of SCM1 (**Figure 2E** and **2F**).

Moreover, to determine the predominant mode of SCM1 cell division, we conducted an intense measurement to observe changes in the diameter and length of SCM1 in the late exponential phase. The results revealed two significant decreases in the length of SCM1 within a 48-hour period, occurring at 11 PM on the Day 6 and 7 AM on the Day 7, respectively (**Figure 2G**). However, no notable change in diameter was observed within the 48-hour timeframe. The time interval between the two decreasing points aligns with the doubling time of SCM1 (Td ≈ 30 h, **Supplementary Figure 2**), indicating a high level of synchronicity in cell division.

### Characterization of intracellular storage granules

Our cryo-ET results revealed density regions corresponding to granule in the cytoplasm of SCM1 (**Figure 1D** and **1E**). Based on differences in electronic density, we classified them into medium-density regions (MDR) and high-density regions (HDR). The MDR observed in SCM1 displayed a distinct and consistent spherical morphology, with diameters typically ranging from approximately 85−185 nm, with a smooth and continuous surface. The HDR exhibited an irregular shape and comprised varying electronic density. Most HDRs were situated at the edge of cell, near the cell membrane (**Figure 1** and **Figure 2**).

To analyze the elemental composition of MDR and HDR granules in the SCM1 cells, we used a Talos F200X to obtain EDS spectra. Area scans revealed differential compositions of the granules, while HDRs containing elevated concentrations of calcium, phosphorus, oxygen, and magnesium, the MDRs had a slight increase in carbon concentration and a slightly lower concentration of oxygen (**Figure 3** and **Supplementary Table 2**). Carbon and nitrogen were the most abundant elements throughout the entire scanned region across the cells.

**Figure 3.**
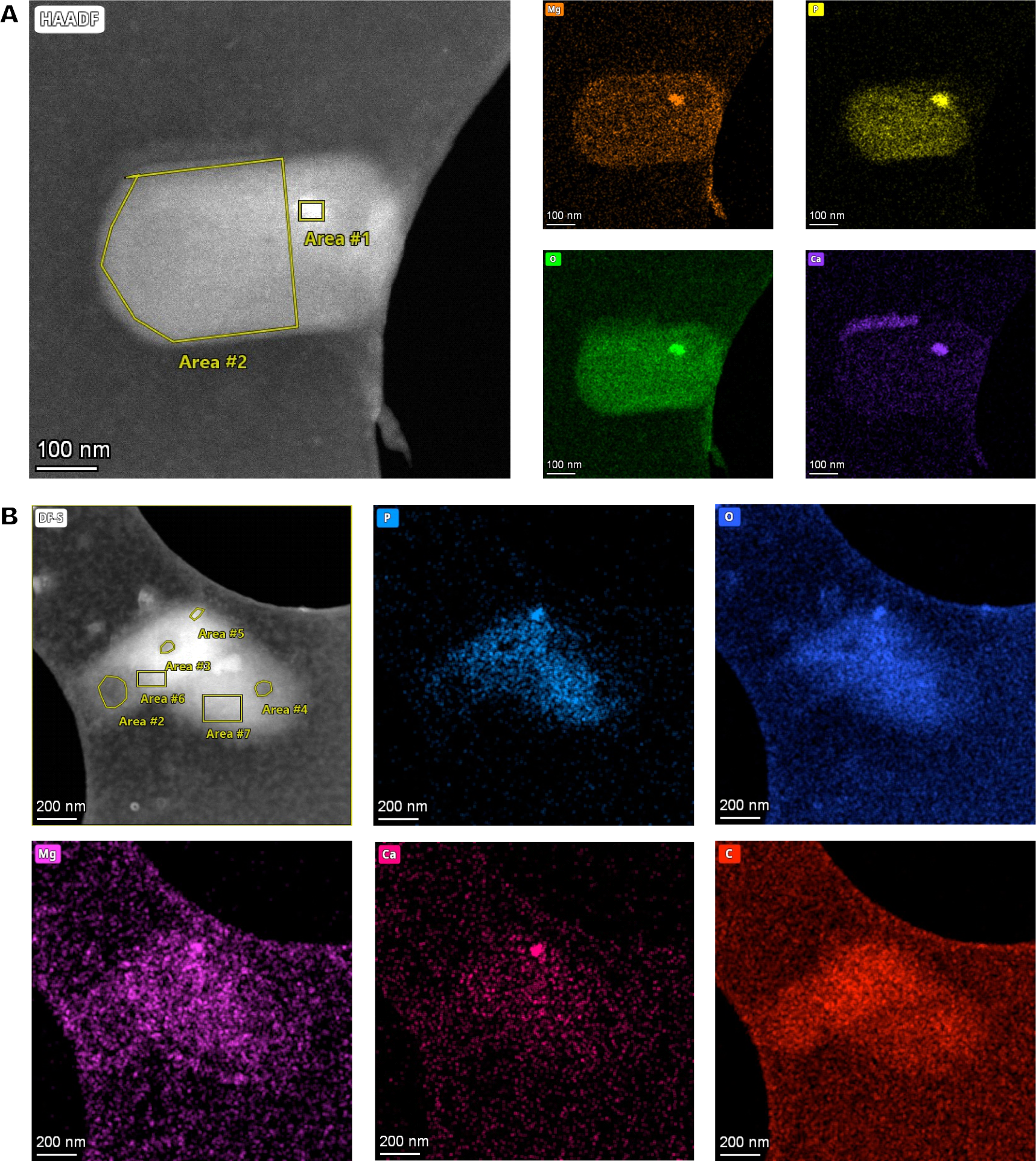
Elements in storage granules of SCM1. (A) STEM images and EDS elemental analysis revealed that magnesium (Mg), phosphorus (P), oxygen (O), and calcium (Ca) in the Area #1 stood out from the background, indicating that enrichment of these elements in the HDR. (B) STEM-EDS results of two adjacent SCM1 cells. Area #5 is a HDR that enriched with Mg, P, O, and Ca. Area #2, 3, and 4 are MDRs with slightly higher concentration of carbon (C).

The mean Raman spectrum of SCM1 showed main biomolecules in single cells (**Figure 4A**). The Raman band at 1726 cm^−1^, which was slightly shifted from 1739 cm^−1^ in *Nitrososphaera gargensis* (Spang et al., 2012), could be attributed to polyhydroxyalkanoate (PHA). This shift of band was caused by the stretching of the C=O ester in PHA (Izumi and Temperini, 2010). The presence of this band also suggested that the PHA produced by SCM1 may have a high crystallinity (Izumi and Temperini, 2010), which was consistent with the results of cryo-ET (**Figure 1**).

**Figure 4.**
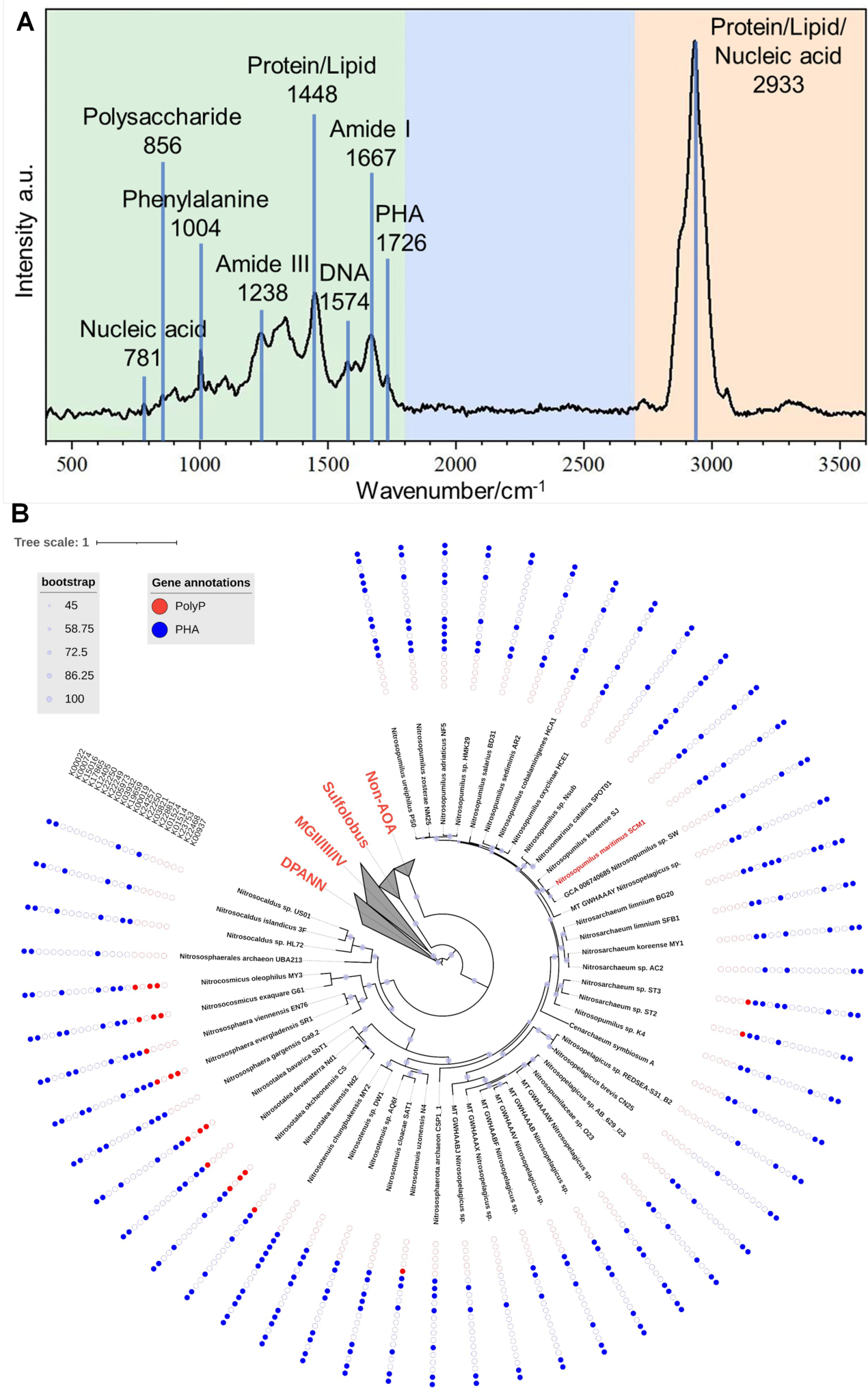
The Raman spectrum of *Nitrosopumilus maritimus* SCM1 and the distributions of metabolic genes regarding PolyP and PHA in AOA. (A) The mean Raman Spectrum of 33 single cells of SCM1. A characteristic peak of PHA at intensity = 1726 cm^−1^ was identified. (B) The phylogenomic tree of 51 nonredundant AOA genomes. The presence or absence of metabolic genes related to PolyP and PHA annotated by KEGG are shown around the tree. Solid dots: presence of the gene, empty dots: absence of the gene.

To further assess SCM1’s potential to metabolize storage granules and fully understand the metabolic potential at higher taxa, we conducted an in-depth exploration of key metabolic pathways in *Nitrosopumilus maritimus* SCM1 and other AOA genomes, focusing on Polyphosphate (PolyP) and polyhydroxyalkanoate (PHA) metabolism. After conducting a thorough genome annotation, we identified several crucial PHA metabolic genes, such as *PhaE* and *PhaC*, which are responsible for PHA synthesis. Additionally, we discovered *BdhA*, along with various hydratases and dehydrogenases, which play a role in PHA degradation in SCM1. However, despite the presence of intracellular PolyP confirmed by STEM-EDS analysis, we found no homologous genes associated with PolyP synthesis and degradation, suggesting additional undiscovered genes involved in PolyP metabolism in SCM1.

When considering a broader spectrum of AOA, we observed the presence of PolyP and PHA metabolic genes across various strains (**Figure 4B**). Notably, PolyP genes were identified in AOA strains ST2, ST3, N4, Nd1, Nd2, CS, SbT1, SR1, EN76, G61, and MY3. Among these strains, only ST2 and ST3 were found in marine habitats, specifically wastewater treatment beside the ocean. The occurrence of *Ppk2* genes in terrestrial AOA strains (CS, SbT1, SR1, G61, MY3) highlighted the ecological diversity and metabolic versatility within the AOA clade.

## Discussion

According to previous studies, SCM1 owns two cell division systems in the genome, but only uses one system, the Cdv system, correlated with the eukaryotic ESCRT-III machinery, for cell division (Pelve et al., 2011; van Wolferen et al., 2022). However, the function of FtsZ system in SCM1 even in the whole *Nitrososphaerota* phylum, is not yet fully understood. Only one study has reported that the absence of polymerization function prevents FtsZ from participating in the cell division process (Ng et al., 2013). Based on the cryo-EM images we captured, we not only observed SCM1 cells in the fission state but also identified cells potentially in the budding state (**Figure 2E** and **2F**), suggesting the activation of the FtsZ system during the budding process. However, given the limited number of SCM1 cells we observed in the budding state, we cannot rule out the possibility of them being extracellular membrane vesicles, similar to those found in haloarchaea *Haloferax volcanii* (Ring and Eichler, 2001; Shalev et al., 2017; Tamir and Eichler, 2017).

Nutrition storage is crucial for microorganisms, providing resilience against fluctuations in nutrient availability. In the previous studies involving bacteria and archaea, similar storage granules had been analyzed using both electron energy loss spectroscopy (EELS) and energy dispersive X-ray spectroscopy (EDS) to determine their elemental composition (Lechaire et al., 2002; Remonsellez et al., 2006). In this study, STEM-EDS analysis has unveiled the intricate elemental composition of intracellular storage granules within *Nitrosopumilus maritimus* SCM1. The HDR granules exhibit heightened levels of phosphate, oxygen, and metals, suggesting their potential involvement in the resilience of oligotrophic marine environments. The discovery of these storage granules opens intriguing avenues for unraveling SCM1’s metabolic strategies and adaptive mechanisms.

In the early 20^th^ century, ultrastructural investigations have already revealed the presence of polyphosphates granules in archaeal cells (Rudnick et al., 1990; Scherer and Bochem, 1983). Toso et al., introduced EDS analysis and demonstrated the high abundance of phosphorus, oxygen and cationic elements in polyphosphate bodies in *Methanospirillum hungatei* (Toso et al., 2011), *Archaeoglobus fulgidus* (Toso et al., 2016) and ANME-2b (McGlynn et al., 2018). Consistent to previous studies, the high concentration of phosphorus and oxygen, along with cationic elements, suggests the presence of polyphosphate (PolyP) in the HDR granules, which indicated that SCM1 may have a greater contribution to global phosphorus cycling than previously recognized (Qin et al., 2020). Divalent cations, such as magnesium and calcium, were known to form ionic bonds between phosphate groups, enabling tighter packing and increased granule density (Parsons et al., 2010). In previous studies, it was believed that carboxysomes, glycogen, and polyphosphate were absent in SCM1 (Urakawa et al., 2011). Instead, an amorphous region with higher electron density, possibly containing phosphorus, was observed. However, our findings from cryo-ET and STEM-EDS analysis have confirmed the presence of polyphosphate in SCM1. This significant discovery was facilitated by advancements in technology and a heightened research focus on understanding the elemental composition of archaea cells.

The presence of PHA in SCM1, as indicated by STEM-EDS, Raman spectroscopy, genome and proteome analyses, opens avenues for understanding its physiological functions. While the exact function of PHA in SCM1 remains to be elucidate (Qin et al., 2020), its potential role in carbon and energy storage, similar to its function in haloarchaea, warrants further investigation. The existence of PHA may contribute to SCM1’s capacity to adapt to severe environmental conditions or insufficient nutrient availability. The 3-hydroxypropionate/4-hydroxybutyrate (3HP/4HB) cycle, discovered in AOA, represents the most energy-efficient aerobic carbon fixation pathway currently known (Johnson et al., 2023; Konneke et al., 2014). According to the definition of the PHA synthesis/degradation pathway, Konneke et al. found a key step involving the synthesis of PHA precursors in the carbon cycle pathway of SCM1 (Konneke et al., 2014).

Our analysis of the PHA synthesis/degradation pathway, considering the presence of key genes like *PhaEC* in the genome and proteome (**Figure 4B** and **Supplementary Table 3**), suggests that under conditions of limited nutrition, such as depletion of bicarbonate ions, the 3HP/4HB cycle in SCM1 may not function normally. Instead, SCM1 could switch to the biosynthesis pathway of PHA, utilizing carbon for survival. This adds credence to the hypothesis of PHA’s presence in SCM1 cells. Further exploration of the role of PHA in SCM1’s metabolism promises valuable insights into its adaptive strategies.

## Materials and Methods

### Nitrosopumilus maritimus SCM1 cultivation

We followed previous protocol for SCM1 cultivation, growth monitoring, and culture purity assessment (Law et al., 2021b). To ensure the absence of bacterial and organic contaminants during the long-term cultivation, we used a combination of 0.5 μg/ml Rifamycin sodium and 1 μg/ml hydrosoluble Amphotericin B. We assessed cell growth by detecting nitrite production, which was determined by diazo-colorimetric assay with photometric detection at 545 nm. For evaluating culture purity, we employed quantitative real-time PCR (qPCR) to accurately measure the culture’s integrity and absence of contaminants, detailed data are provided in **Supplementary Table 1**.

### Collection and processing of Raman spectra of Nitrosopumilus maritimus SCM1

*Nitrosopumilus maritimus* SCM1 was incubated at 30°C for 14 days. The pre-treatment processes were as described previously (Wang et al., 2021). Single-cell Raman spectra were acquired using a Raman imaging microscope (WITec alpha 300R) with 532 nm laser. All raw spectra were preprocessed with Project Five 5.2 (WITec) for baseline correction and vector normalization.

### Cryo-EM sample preparation and data acquisition

Quantifoil Cu R2/1 grids were glow discharged at 15 mA for 45 seconds using a Pelco easiGlow discharged unit. An aliquot of 4 μL concentrated sample of SCM1 was applied to the front side of the grids, then the sample was vitrified by plunge freezing in a liquid ethane using a Mark IV Vitrobot (Thermo Fisher Scientific) at 6 °C and 100% humidity. The vitrified samples were loaded into a 300-kV Titan Krios microscopy G3i (Thermo Fisher Scientific) equipped with GIF quantum energy filter and K2 Summit direct electron detector (Gatan Inc.). Images were acquired using SerialEM package (Mastronarde, 2003) by the K2 camera operated under the super-resolution mode at a nominal magnification of ×26,000 (calibrated pixel size 5.417 Å). Each micrograph was fractionated into 36 frames with a total dose of 50 e^−^/Å^2^. The defocus range was set from −5.0 to −8.0 μm.

### Cryo-ET data collection and tomogram reconstruction

For cryo-ET, the vitrified samples were loaded into a 300-kV Titan Krios microscopy G4 (Thermo Fisher Scientific) equipped with Selectris energy filter and CEOS CCOR-spherical aberration / chromatic aberration (Cs/Cc) corrector. Tilt images were acquired by Falcon 4 camera under counting mode at a nominal magnification of ×26,000 (calibrated pixel size of 3.41 Å). A dose-symmetric scheme (Hagen et al., 2017) was used to collect tilt-series from −60° to +60° at a step size of 2° using SerialEM software (Mastronarde, 2003). Each tilt image was recorded as a video stack consisting of 10 frames with a dose of 1.64 e^−^/Å^2^. The defocus was set range from −4 to −6 μm and the total dose was 100 e^−^/Å^2^. Video frames of each tilt image were motion-corrected by MotionCor2 v.1.1.0 (Zheng et al., 2017). Tilt series were merged into one stack and aligned by patch tracking in IMOD v.4.9.12 (Kremer et al., 1996). In total, 221 sets of tilt-series were collected. The selected tomograms were reconstructed the missing-wedge information and increased signal-to-noise ratio by IsoNet (Liu et al., 2022) for display and segmentation by Amira (version 2020.3.1, Thermo Fisher Scientific).

### STEM-EDS analysis

Samples were cryo-transferred by the Elsa™ holder (Gatan, model 698, ultra-low profile) and imaged in a Thermofisher Talos F200X transmission electron microscopy equipped with a Ceta camera in bright field mode, operated at a 200 kV acceleration voltage. Next, the cryo-holder with the sample was inserted into a pumping station to warm up to room temperature. Then a focused electron probe with high energy was used to stimulate a specific area of the sample. The atoms being stimulated may eject electrons, resulting in the emission of X-ray photons as a form of energy release. The energy exhibited by these X-rays is indicative of the particular atomic element from which they originated. Therefore, an energy-dispersive spectrometer can be employed to quantify the quantity and energy of the X-rays, providing a quantitative description of the elemental composition of the stimulated area of the sample (Toso et al., 2011). STEM-EDS tomography was acquired at room temperature. The data process and 3D visualization were performed by Inspect 3D and Avizo software.

### Gene and genomic analysis

Phylogenomic analysis of *Nitrososphaerota* was conducted using IQ-TREE with GTDB archaeal markers obtained from the Genome Taxonomy Database. The selected markers were carefully aligned using MAFFT and then concatenated into a supermatrix. The best-fitting substitution model was determined using ModelFinder, and the phylogenomic tree was constructed using Maximum Likelihood with ultrafast bootstrap approximation. The genomes were curated as our previous study. The PolyP and PHA were investigated in AOA after predicting proteins from the AOA genomes. The obtained AOA proteins were aligned against the KO profiles available in the KEGG database (accessed in November 2023). The results were filtered based on the HMM score (>90), and the corresponding KO related to PolyP and PHA metabolic genes were searched. Additionally, to confirm our annotation, the presence of each key functional domain of the targeted proteins was manually checked using InterProScan (https://www.ebi.ac.uk/interpro/). The KO numbers of PolyP related genes are: K00937(*Ppk1*), K22468(*Ppk2*), K23753(*Ppk2*), K01514(*PPX1*), K01524(*Ppx/GppA*), and PHA related genes are: K22881(*PhaE*), K03821(*PhaC*), K22250(*PhaB*), K24257(*PhaR*), K00019(*BdhA*), K19659(*PhaJ*), K03932(*PhaZ1*), K05973(*PhaZ*), K22249(*PhaZ*), K22250(*PhaZ*), K12405(hydratase), K17865(hydratase), K15016(hydratase), K00074(dehrogenase), K00022(dehrogenase). The resulting tree was beautified using the Interactive Tree of Life (iTOL) web tool. The Newick-formatted tree was uploaded to iTOL, where branches were colorized, additional annotations were added, and the overall appearance of the tree was customized for better clarity and aesthetics.

### Proteome analysis

Filters containing late exponential phase *Nitrosopumilus maritimus* SCM1 cells were extracted using a modified protocol as described before (Qin et al., 2018), either trypsin or GluC was used as protease digestion. Mass spectrometry analysis was carried out at Shenzhen Wininnovate BioTechnology Co., Ltd. DDA (data-dependent acquisition) mass spectrum techniques were used to acquire tandem MS data on a ThermoFisher Q Exactive mass spectrometer fitted with a Nano Flex ion source. The raw MS/MS spectra were treated and searched against the NCBI and UniProt Proteome dataset using MaxQuant 2.0.3.0. The generated data set contains the results of different hydrolases and databases for comparison.

## Supporting information

Supplementary Tabel 3

## Acknowledgements

We thank Huiqin Xu, Xiaoyun Yang, and Zongqiang Li for assistance on experiment design. Jing Wu, Yuanzhu Gao, Juying Tan, and Fangfang Zhang for technical support. This study was supported by the Stable Support Plan Program of Shenzhen Natural Science Fund (20200925173954005), National Natural Science Foundation of China (32241028 and 91851210), the Shenzhen Key Laboratory of Marine Archaea Geo-Omics, Southern University of Science and Technology (ZDSYS201802081843490), and the Project of Educational Commission of Guangdong Province of China (2020KTSCX123). The bioinformatic analyses were performed on the Tai-Yi high-performance supercomputer cluster at Southern University of Science and Technology.

## Author contributions

ZL, CZ designed the study. YZ, AY, JY, WH performed the sample preparation, data analyses, and wrote the manuscript. SG conducted the reconstruction of cryo-ET data. YL prepared cell cultures, performed the sample preparation experiment. YD, XP performed the STEM-EDS analyses. DC performed the Raman spectrum analyses. OP, WT, RB, SC, LF assisted the lab work or data analyses. PW, SS provided platforms. WQ, YL assisted study design, data interpretation and manuscript editing. ZL, CZ supervised the investigation and revised manuscript.

