## Supplementary Information for "The 3D ultrastructural morphology of a marine ammonia-oxidizing archaeon"

Supplementary Figure 1. Growth curve of *Nitrosopumilus maritimus* SCM1 with 1mM of primary $NH_4$ concentration.

Supplementary Figure 2. Statistically generated chart depicting the size range of SCM1 cells.

Supplementary Figure 3. Images of SCM1 at different degrees by traditional Cryo-EM.

Supplementary Table 1. The purity of the SCM1 cultures was evaluated using the qPCR.

Supplementary Table 2. The STEM-EDS data of selected *Nitrosopumilus maritimus* SCM1 cells.

Supplementary Table 3. The proteome data of *Nitrosopumilus maritimus* SCM1.

Supplementary Movie 1. Reconstructed 3D cryo-tomogram of a typical cell of *Nitrosopumilus maritimus* SCM1.

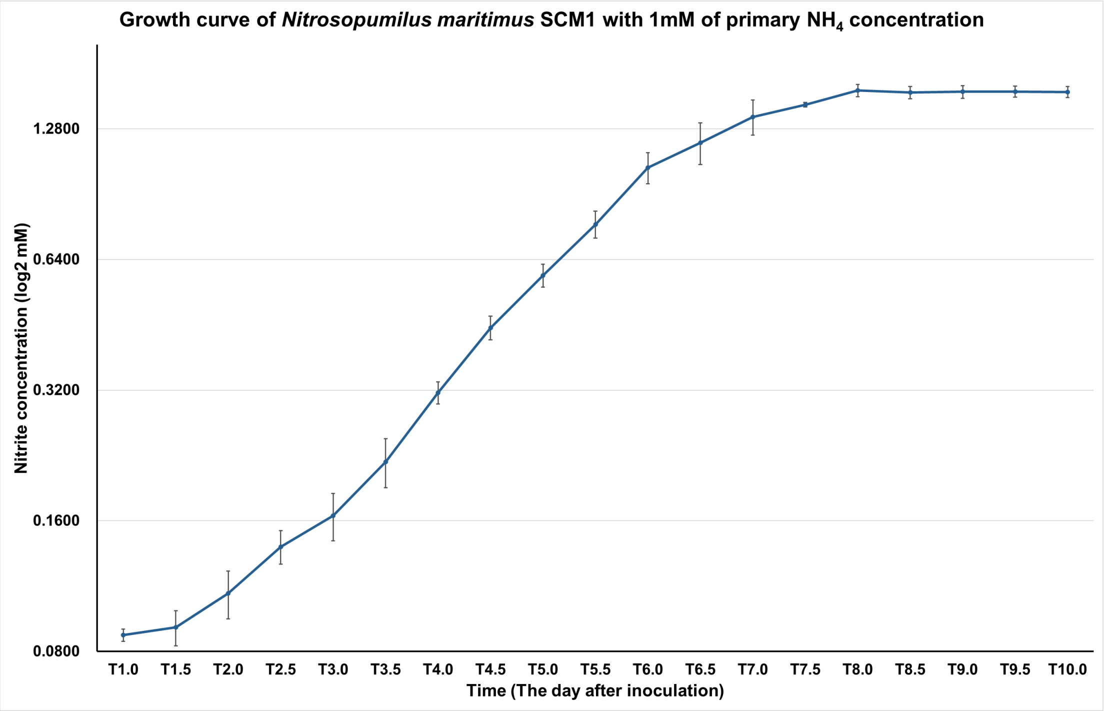


**Supplementary Figure 1. Growth curve of *Nitrosopumilus maritimus* SCM1 with 1mM of primary $NH_4$ concentration.**

We monitored cell growth in *Nitrosopumilus maritimus* SCM1 cultures by analyzing nitrite concentration. Nitrite concentration was continued recorded for every 12 hours after inoculation. The growth curve was plotted from 6 replicates with means ± SDs (error bars).

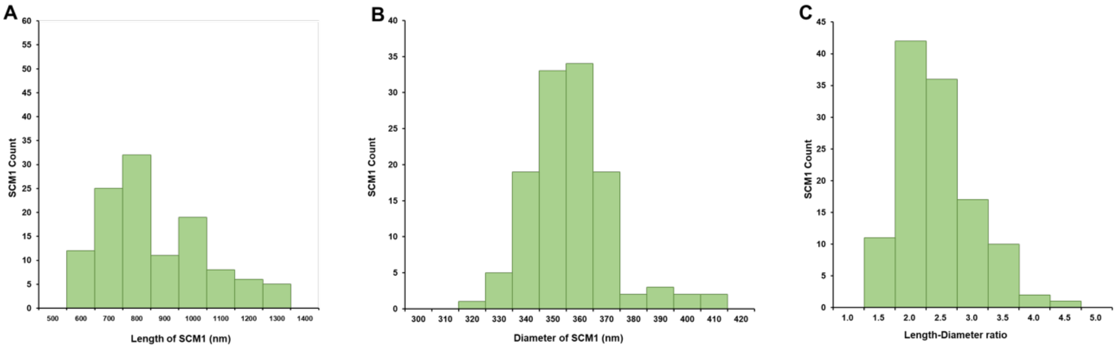


**Supplementary Figure 2. Statistically generated chart depicting the size range of SCM1 cells.**

The y-axis represents the number of SCM1 cells and the x-axis in each panel represents (A) Length (nm), (B) Diameter (nm), (C) Lenth-Diameter ratio of SCM1. The average length was 860 ± 181 nm (maximum: 1349 nm, minimum: 593 nm). The average diameter was 357 ± 15 nm (maximum: 412 nm, minimum 324 nm). The mean length-diameter ratio was 2.43 ± 0.57.

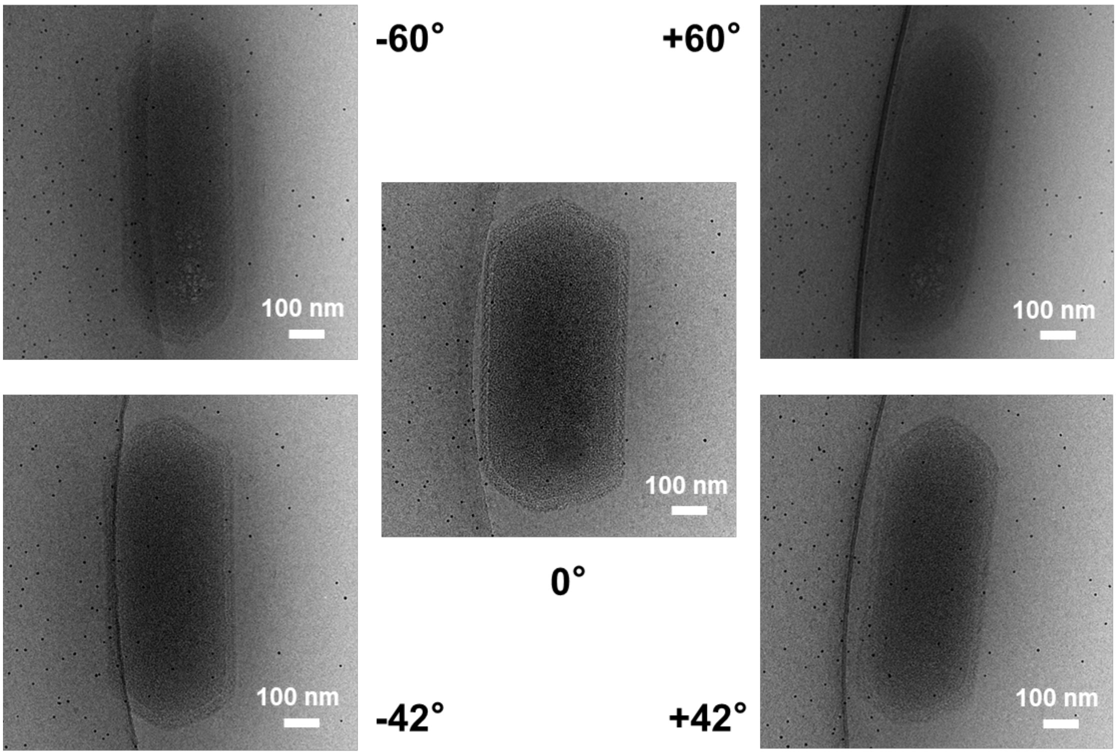


**Supplementary Figure 3. Images of SCM1 at different degrees by traditional Cryo-EM.**

Images of SCM1 at different angles were acquired by traditional cryo-EM. It can be found that the structural details of SCM1 at high-degree angle such as ±42° or ±60° became unclear and the contrast was relatively low.

**Supplementary Table 1. The purity of the SCM1 cultures was evaluated using the qPCR.**

| Sample | Mean copies of Bacteria | Mean copies of Archaea | Percentage of SCM1(%) |
|---|---|---|---|
| SCM1-1-220716 | 3,320.15 | 101,518,853.33 | 100.00 |
| SCM1-1-220801 | 2,822.03 | 127,649,797.33 | 100.00 |
| SCM1-1-220915 | 4,408.67 | 225,270,208.00 | 100.00 |
| SCM1-1-221110 | 3,348.08 | 136,850,829.33 | 100.00 |
| SCM1-1-221124 | 2225.04 | 126,674,933.33 | 100.00 |
| SCM1-1-221219 | 591.78 | 240,940,053.33 | 100.00 |
| SCM1-1-230212 | 746.39 | 63,871,446.00 | 100.00 |
| SCM1-1-230424 | 12,914.45 | 719,556,885.33 | 100.00 |
| SCM1-1-230515 | 9,383.56 | 212,321,997.33 | 100.00 |
| SCM1-1-231009 | 1,350.29 | 101,788,965.33 | 100.00 |

**Supplementary Table 2. The STEM-EDS data of selected *Nitrosopumilus maritimus* SCM1 cells.**

In Figure 3A, two areas were emphasized:

| Area 1 | Z | Element | Family | Atomic Fraction (%) | Atomic Error (%) | Mass Fraction (%) | Mass Error (%) | Fit Error (%) |
|---|---|---|---|---|---|---|---|---|
| | 6 | C | K | 56.37 | 2.76 | 44.71 | 2.4 | 3.1 |
| | 7 | N | K | 11.62 | 2.12 | 10.75 | 1.99 | 1.12 |
| | 8 | O | K | 19.74 | 3.23 | 20.86 | 3.37 | 0.62 |
| | 11 | Na | K | 3.57 | 0.71 | 5.43 | 1.07 | 1.98 |
| | 12 | Mg | K | 2.02 | 0.41 | 3.24 | 0.65 | 2.59 |
| | 13 | Al | K | 0.12 | 0.04 | 0.22 | 0.07 | 24.03 |
| | 14 | Si | K | 0.19 | 0.05 | 0.35 | 0.09 | 17.18 |
| | 15 | P | K | 3.27 | 0.63 | 6.7 | 1.24 | 1.36 |
| | 16 | S | K | 0.48 | 0.1 | 1.01 | 0.21 | 7.27 |
| | 17 | Cl | K | 0.75 | 0.15 | 1.75 | 0.34 | 5.2 |
| | 20 | Ca | K | 1.81 | 0.26 | 4.8 | 0.66 | 1.69 |
| | 26 | Fe | K | 0.05 | 0.02 | 0.19 | 0.07 | 33.87 |

| Area 2 | Z | Element | Family | Atomic Fraction (%) | Atomic Error (%) | Mass Fraction (%) | Mass Error (%) | Fit Error (%) |
|---|---|---|---|---|---|---|---|---|
| | 6 | C | K | 67.53 | 2.53 | 58.89 | 2.48 | 1.42 |
| | 7 | N | K | 14.51 | 2.51 | 14.75 | 2.56 | 0.3 |
| | 8 | O | K | 10.52 | 1.92 | 12.22 | 2.19 | 1.37 |
| | 11 | Na | K | 4.68 | 0.91 | 7.82 | 1.47 | 0.25 |
| | 12 | Mg | K | 0.59 | 0.12 | 1.04 | 0.21 | 0.54 |
| | 13 | Al | K | 0.04 | 0.01 | 0.07 | 0.02 | 3.42 |
| | 14 | Si | K | 0.07 | 0.01 | 0.15 | 0.03 | 2.7 |
| | 15 | P | K | 0.97 | 0.19 | 2.17 | 0.41 | 0.31 |
| | 16 | S | K | 0.32 | 0.06 | 0.74 | 0.14 | 0.39 |
| | 17 | Cl | K | 0.48 | 0.09 | 1.24 | 0.23 | 0.36 |
| | 20 | Ca | K | 0.25 | 0.03 | 0.72 | 0.1 | 0.53 |
| | 26 | Fe | K | 0.05 | 0.01 | 0.18 | 0.03 | 1.39 |

In Figure 3B, six areas were emphasized:

| Area 2 | Z | Element | Family | Atomic Fraction (%) | Atomic Error (%) | Mass Fraction (%) | Mass Error (%) | Fit Error (%) |
|---|---|---|---|---|---|---|---|---|
| | 6 | C | K | 73.37 | 2.77 | 63.59 | 3.12 | 10.75 |
| | 7 | N | K | 7.07 | 1.48 | 7.15 | 1.47 | 5.43 |
| | 8 | O | K | 7.93 | 1.6 | 9.16 | 1.8 | 2 |
| | 9 | F | K | 4.69 | 1 | 6.43 | 1.33 | 5.35 |
| | 11 | Na | K | 2.56 | 0.55 | 4.25 | 0.88 | 3.41 |
| | 12 | Mg | K | 1.25 | 0.27 | 2.19 | 0.47 | 4.74 |
| | 13 | Al | K | 0.2 | 0.07 | 0.4 | 0.13 | 24.92 |
| | 14 | Si | K | 1.46 | 0.31 | 2.97 | 0.61 | 4.61 |
| | 15 | P | K | 0.27 | 0.08 | 0.61 | 0.17 | 18.36 |
| | 16 | S | K | 0.36 | 0.09 | 0.84 | 0.2 | 13 |
| | 17 | Cl | K | 0.32 | 0.08 | 0.81 | 0.2 | 14.21 |
| | 19 | K | K | 0.11 | 0.04 | 0.31 | 0.13 | 35.4 |
| | 20 | Ca | K | 0.26 | 0.05 | 0.74 | 0.14 | 11.38 |
| | 26 | Fe | K | 0.14 | 0.04 | 0.55 | 0.14 | 21.58 |

| Area 3 | Z | Element | Family | Atomic Fraction (%) | Atomic Error (%) | Mass Fraction (%) | Mass Error (%) | Fit Error (%) |
|---|---|---|---|---|---|---|---|---|
| | 6 | C | K | 67.85 | 5.21 | 57.43 | 5.75 | 21.65 |
| | 7 | N | K | 8.01 | 1.99 | 7.9 | 1.87 | 7.99 |
| | 8 | O | K | 11.05 | 2.61 | 12.46 | 2.76 | 5.04 |
| | 9 | F | K | 5.32 | 1.42 | 7.12 | 1.8 | 12.07 |
| | 11 | Na | K | 2.96 | 0.79 | 4.8 | 1.21 | 10.79 |
| | 12 | Mg | K | 1.04 | 0.34 | 1.79 | 0.56 | 20.98 |
| | 13 | Al | K | 0.35 | 0.21 | 0.67 | 0.4 | 55.49 |
| | 14 | Si | K | 1.26 | 0.37 | 2.5 | 0.7 | 16.93 |
| | 15 | P | K | 0.55 | 0.23 | 1.21 | 0.49 | 33.37 |
| | 16 | S | K | 0.45 | 0.2 | 1.03 | 0.45 | 37.49 |
| | 17 | Cl | K | 0.66 | 0.23 | 1.65 | 0.54 | 24.84 |
| | 19 | K | K | 0.11 | 0.16 | 0.3 | 0.43 | 141.63 |
| | 20 | Ca | K | 0.3 | 0.13 | 0.86 | 0.36 | 37.37 |
| | 26 | Fe | K | 0.07 | 0.12 | 0.28 | 0.46 | 162.32 |

| Area 4 | Z | Element | Family | Atomic Fraction (%) | Atomic Error (%) | Mass Fraction (%) | Mass Error (%) | Fit Error (%) |
|---|---|---|---|---|---|---|---|---|
| | 6 | C | K | 71.77 | 3.1 | 61.61 | 3.45 | 11.29 |
| | 7 | N | K | 5.83 | 1.28 | 5.84 | 1.26 | 7.77 |
| | 8 | O | K | 11.71 | 2.31 | 13.39 | 2.55 | 3.27 |
| | 9 | F | K | 3.7 | 0.92 | 5.03 | 1.21 | 13.45 |
| | 11 | Na | K | 2.19 | 0.52 | 3.61 | 0.83 | 10.24 |
| | 12 | Mg | K | 0.9 | 0.25 | 1.57 | 0.43 | 17.68 |
| | 13 | Al | K | 0.27 | 0.15 | 0.52 | 0.28 | 50.02 |

| | | | | | | | |
|---|---|---|---|---|---|---|---|
| 14 | Si | K | 1.78 | 0.4 | 3.58 | 0.78 | 8.69 |
| 15 | P | K | 0.31 | 0.14 | 0.68 | 0.31 | 40.95 |
| 16 | S | K | 0.34 | 0.14 | 0.79 | 0.32 | 35.65 |
| 17 | Cl | K | 0.74 | 0.19 | 1.88 | 0.48 | 16.65 |
| 19 | K | K | 0.12 | 0.11 | 0.32 | 0.31 | 93.69 |
| 20 | Ca | K | 0.12 | 0.08 | 0.35 | 0.23 | 64.52 |
| 26 | Fe | K | 0.21 | 0.09 | 0.84 | 0.35 | 39.33 |

| Area 5 | Z | Element | Family | Atomic Fraction (%) | Atomic Error (%) | Mass Fraction (%) | Mass Error (%) | Fit Error (%) |
|---|---|---|---|---|---|---|---|---|
| | 6 | C | K | 35.45 | 5.65 | 24.51 | 4.5 | 22.88 |
| | 7 | N | K | 7.55 | 1.69 | 6.09 | 1.34 | 8.58 |
| | 8 | O | K | 23.22 | 4.15 | 21.38 | 3.72 | 2.67 |
| | 9 | F | K | 12.44 | 2.58 | 13.61 | 2.68 | 6.1 |
| | 11 | Na | K | 7.89 | 1.69 | 10.45 | 2.1 | 4.6 |
| | 12 | Mg | K | 3.94 | 0.9 | 5.51 | 1.2 | 7.41 |
| | 13 | Al | K | 0.16 | 0.2 | 0.25 | 0.31 | 126.6 |
| | 14 | Si | K | 1.26 | 0.34 | 2.04 | 0.54 | 16.84 |
| | 15 | P | K | 4.05 | 0.87 | 7.22 | 1.46 | 5.78 |
| | 16 | S | K | 0.65 | 0.23 | 1.2 | 0.41 | 27.97 |
| | 17 | Cl | K | 0.7 | 0.23 | 1.42 | 0.45 | 24.96 |
| | 19 | K | K | 0.04 | 0.16 | 0.1 | 0.35 | 350.46 |
| | 20 | Ca | K | 2.54 | 0.44 | 5.85 | 0.92 | 5.2 |
| | 26 | Fe | K | 0.12 | 0.12 | 0.37 | 0.39 | 103.72 |

| Area 6 | Z | Element | Family | Atomic Fraction (%) | Atomic Error (%) | Mass Fraction (%) | Mass Error (%) | Fit Error (%) |
|---|---|---|---|---|---|---|---|---|
| | 6 | C | K | 53.89 | 3.69 | 43 | 3.45 | 11.43 |
| | 7 | N | K | 15.33 | 2.82 | 14.27 | 2.61 | 2.12 |
| | 8 | O | K | 12.66 | 2.39 | 13.46 | 2.48 | 1.56 |
| | 9 | F | K | 6.62 | 1.36 | 8.36 | 1.66 | 4.44 |
| | 11 | Na | K | 4.71 | 0.97 | 7.2 | 1.43 | 2.43 |
| | 12 | Mg | K | 1.7 | 0.37 | 2.75 | 0.58 | 4.75 |
| | 13 | Al | K | 0.31 | 0.08 | 0.55 | 0.14 | 16.5 |
| | 14 | Si | K | 1.52 | 0.32 | 2.84 | 0.58 | 4.63 |
| | 15 | P | K | 0.75 | 0.16 | 1.54 | 0.33 | 7.61 |
| | 16 | S | K | 0.71 | 0.15 | 1.52 | 0.32 | 7.52 |
| | 17 | Cl | K | 1.29 | 0.26 | 3.04 | 0.59 | 4.83 |
| | 19 | K | K | 0.12 | 0.05 | 0.31 | 0.12 | 33.97 |
| | 20 | Ca | K | 0.25 | 0.05 | 0.66 | 0.13 | 12.57 |
| | 26 | Fe | K | 0.14 | 0.04 | 0.52 | 0.14 | 21.51 |

| Area 7 | Z | Element | Family | Atomic Fraction (%) | Atomic Error (%) | Mass Fraction (%) | Mass Error (%) | Fit Error (%) |
|---|---|---|---|---|---|---|---|---|

| | | | | | | | |
|---|---|---|---|---|---|---|---|
| 6 | C | K | 50.81 | 3.87 | 39.46 | 3.53 | 12.18 |
| 7 | N | K | 10.03 | 1.97 | 9.08 | 1.78 | 2.72 |
| 8 | O | K | 18.91 | 3.32 | 19.56 | 3.34 | 0.59 |
| 9 | F | K | 7.47 | 1.5 | 9.18 | 1.79 | 2.04 |
| 11 | Na | K | 4.8 | 0.99 | 7.13 | 1.42 | 1.54 |
| 12 | Mg | K | 2.03 | 0.43 | 3.19 | 0.66 | 1.91 |
| 13 | Al | K | 0.29 | 0.07 | 0.5 | 0.12 | 12.48 |
| 14 | Si | K | 1.5 | 0.31 | 2.73 | 0.55 | 2.43 |
| 15 | P | K | 1.51 | 0.31 | 3.02 | 0.6 | 3.29 |
| 16 | S | K | 0.61 | 0.13 | 1.27 | 0.26 | 4.43 |
| 17 | Cl | K | 1.53 | 0.3 | 3.52 | 0.67 | 2.09 |
| 19 | K | K | 0.17 | 0.04 | 0.42 | 0.09 | 12.04 |
| 20 | Ca | K | 0.29 | 0.05 | 0.74 | 0.12 | 5.46 |
| 26 | Fe | K | 0.05 | 0.02 | 0.19 | 0.06 | 28.7 |

**Supplementary Table 3. The proteome data of *Nitrosopumilus maritimus* SCM1.** See separate file in the supplemental data.
